# A highly stable, nondigestible lectin from *Pomacea diffusa* unveils clade-related protection systems in apple snail eggs

**DOI:** 10.1101/2020.06.23.167262

**Authors:** TR Brola, MS Dreon, JW Qiu, H Heras

**Author notes:** Corresponding author: Marcos S. Dreon, Instituto de Investigaciones Bioquímicas de La Plata (INIBIOLP), Consejo Nacional de Investigaciones Científicas y Técnicas (CONICET) - Universidad Nacional de La Plata (UNLP), 60 y 120, 1900 La Plata, Argentina.

## Abstract

The acquisition of egg protection is vital for species survival. Poisonous eggs from *Pomacea* apple snails have defensive macromolecules for protection. Here we isolated and characterized a novel lectin called PdPV1 that is massively accumulated in the eggs of *Pomacea diffusa* and seems part of its protective cocktail. The native protein, an oligomer of ca. 256 kDa, has high structural stability, withstanding 15 min boiling and denaturing by sodium dodecyl sulphate. It resists *in vitro* proteinase digestion and displays structural stability between pH 2.0–12.0 and up to 85 °C. These properties, as well as its subunit sequences, glycosylation pattern, presence of carotenoids, size, and global shape resemble those of its orthologs from other *Pomacea*. Further, like members of the *canaliculata* clade, PdPV1 is recovered unchanged in faeces of mice ingesting it, supporting an antinutritive defensive function. PdPV1 also displays a strong hemagglutinating activity specifically recognizing selected ganglioside motifs with high affinity. This activity is only shared with PsSC, a perivitelline from the same clade (*bridgesii* clade). As a whole, these results indicate that species in the genus *Pomacea* have diversified their eggs defences: Those from the *bridgesii* clade are protected mostly by non-digestible lectins that lower the nutritional value of eggs, in contrast with protection by neurotoxins of other *Pomacea* clades, indicating apple snail egg defensive strategies are clade-specific. The harsh gastrointestinal environment of predators would have favoured their appearance, extending by convergent evolution the presence of plant-like highly stable lectins, a strategy not reported in other animals.

**Summary statement:** Analysis of key snail egg proteins shows evolutionary defensive trends associated with the phylogenetic position, extending by convergent evolution the presence of plant-like defensive strategies not reported in other animals

## Introduction

*Pomacea* freshwater snails (Caenogastropoda, Ampullariidae), particularly those of invasive species belonging to the *canaliculata* clade, lay colourful and poisonous egg masses on emergent substrates above the water level, thus exposing eggs to environmental stressful agents and terrestrial predators (Przeslawski, 2004; Hayes *et al*., 2015). These defences pay off and as a result their eggs have almost no reported predators except for the fire ant *Solenopsis geminata* (Yusa, 2001). We have found that these defences are provided by proteins (perivitellins) already active in the albumen gland of females where the egg components are synthesized (Cadierno *et al*., 2018; Cadierno, Dreon and Heras, 2018), a likely explanation why the brown rat (*Rattus norvegicus*) and the snail kite (*Rostramus sociabilis*) which predate on adult snails, systematically discard the albumen gland (Yusa, Sugiura and Ichinose, 2000).

Extensive studies of the most abundant perivitellins in *Pomacea* species, collectively called PV1s, have shown that they are all oligomeric carotenoproteins of high molecular weight, representing ~70% of the PVF soluble protein (Dreon, Heras and Pollero, 2003; Ituarte *et al*., 2008; Pasquevich, Dreon and Heras, 2014). These carotenoproteins are highly stable in a wide range of pH and temperatures (Dreon, Ceolín and Heras, 2007; Dreon *et al*., 2008; Sun *et al*., 2012; Pasquevich *et al*., 2017) and withstand gastrointestinal digestion (Dreon, Ituarte and Heras, 2010; Ituarte *et al*., 2012; Dreon *et al*., 2013; Pasquevich *et al*., 2017). Nevertheless, some differences emerged: Although the PV1 carotenoprotein of the non-invasive species *P. scalaris* (PsSC) is a lectin (carbohydrate binding protein) with a remarkably strong activity (Ituarte *et al*., 2012, 2018), its orthologues from the invasive species *P. canaliculata* and *P. maculata* (PcOvo and PmPV1, respectively) lack lectin activity.

Many lectins have been experimentally identified in molluscs, mostly in eggs or in organs involved in the synthesis of egg components. Probably, the best-known gastropod egg lectin is *Helix pomatia* agglutinin, which nowadays has important biomedical applications as a histopathological biomarker of tumours displaying differential glycosylation patterns (Sanchez *et al*., 2006). In ampullariids, lectin activity has been reported in the eggs of *Pila ovata* (Uhlenbruck, Steinhausen and Cheesman, 1973) and *Pomacea urceus* (Baldo and Uhlenbruck, 1974) and assumed to be part of the innate immune system against microbial invasions (Prokop, Uhlenbruck and Kohler, 1968). However, the only ampullariid lectin isolated and functionally characterized so far is the PV1 carotenoprotein PsSC from *Pomacea scalaris* eggs. This lectin role immediately suggests an unexpectedly functional divergence within *Pomacea* egg carotenoproteins (Ituarte *et al*., 2018), but this hypothesis needs further comparative analysis. In addition to PV1s, the conspicuous pink-reddish eggs of the invasive species have a neurotoxic perivitelline lethal to mice (Heras *et al*., 2008; Giglio *et al*., 2016) that is absent in the noninvasive species *P. scalaris* which has pale pinkish eggs. This prompted us to study whether the different defensive roles of perivitellins against predation was associated with their phylogeny (Hayes, Cowie and Thiengo, 2009), which clusters *P. maculata* and *P. canaliculata* in themore defived *canaliculata* clade and *P. scalaris* and *P. diffusa* in the more basal *bridgesii* clade.

To better understand carotenoprotein roless and evolutionary trends in *Pomacea* egg defences we studied *Pomacea diffusa*, a member of the basal clade analysing its egg perivitellin fluid and characterizing PdPV1, a novel PV1 carotenoprotein with lectin activity, reporting its structure, structural stability, and functional features in a phylogenetic framework. We suggest a putative role of PdPV1 in embryo protection as both an antinutritive protein (highly stable, nondigestible) and as a toxic lectin. Further, we describe an evolutionary association among *Pomacea* egg defence systems, their phylogenetic position and invasiveness.

## Methods

### Ethics Statement

Studies performed with animals (Protocol Number: P01-01-2016) were approved by the “Comité Institucional para el Cuidado y Uso de Animales de Laboratorio” (CICUAL) of the School of Medicine, Universidad Nacional de La Plata (UNLP) and in accordance with the Guide for the Care and Use of Laboratory Animals.

### Animals

Female BALB/c mice were obtained from the Experimental Animals Lab, School of Veterinary Science, National University of La Plata (UNLP), and weighted around 16 g. Adults of *P. diffusa* were purchased from a local aquarium and genetically identified by sequencing the cytochrome oxidase C subunit I gene, using LCO 1490 and HCO 2198 primers for DNA amplification (Matsukura *et al*., 2013). Sequence was then compared with those deposited in GeneBank (NCBI). Snails were raised in the laboratory in 50 L aquaria, at 25 °C and fed with lettuce and fish food *ad libitum*. Egg masses were collected as soon as they were laid and were never beyond the morula stage.

### Reagents

Reagents used were Carlo Erba Reagents S.A.S or Merck/Sigma chemicals co. Molecular standards and nitrocellulose strips were purchased to GE Healthcare, Uppsala, Sweden. Lectins, streptavidin conjugate and ABC system for electrochemiluminescence were Vector Labs, Burlingame, CA, USA. Goat anti-rabbit horseradish-peroxidase conjugated secondary antibody used was bought to BioRad Laboratories, Inc. Alexa Fluor 488 Protein Labelling kit was Invitrogen, Life Technologies-Molecular Probes. U-shaped microtiter plates for hemagglutinating assays were Greiner Bio One, Germany.

### Protein isolation and purification

Egg homogenate was prepared in ice-cold 20 mM Tris-HCl, pH 7.4 using a Potter type homogenizer. The buffer:sample ratio was kept at 3:1 (v/w). The homogenate was then sequentially centrifuged at 10,000xg for 20 min in an Avanti JE centrifuge (Beckman, Palo Alto, CA) and at 100,000xg for 45 min on a Beckman L8M centrifuge at 10 °C. Both pellets were discarded and the supernatant (henceforth referred to as perivitelline fluid [PVF]) was layered on NaBr δ=1.28 g/mL and centrifuged at 207,000g for 22 h at 10 °C on a Beckman L8M centrifuge (Beckman, Palo Alto, CA). A tube layered with 20 mM Tris-HCl in lieu of the sample was used as a blank for density calculations. Subsequently, aliquots of 200μL were collected sequentially starting at the top of the tube. Absorbance of each aliquot was determined at 280 nm to obtain the protein profile. The refractive index of the blank tube aliquots was determined at 26 °C with a refractometer (Bausch and Lomb, New York, New York) and converted to density using tabulated values (Orr, Adamson and Lindgren, 1991). Fractions showing absorbance peaks at 280 nm were pooled and purified by size exclusion chromatography (SEC) in a HPLC system (Agilent technologies, 1260 infinity, Waldron, Germany) using a Superdex 200 HR 10/300 GL column (GE Healthcare Bio-Sciences AB, Uppsala, Sweden) as described by Ituarte et al (Ituarte *et al*., 2018). Purification process of the protein (hereafter named PdPV1) was checked by polyacrylamide gel electrophoresis (PAGE) as described below. Protein content was determined using BSA as standard (Lowry *et al*., 1951) in an Agilent 8453 UV/Vis diode array spectrophotometer (Agilent Technologies). PsSC was prepared as described elsewhere (Ituarte *et al*., 2008).

### Polyacrylamide gel electrophoresis

Native (non-denaturing) PAGE was performed in a 4–20% gradient gel in a Mini-Protean III System (Bio Rad Laboratories, Inc.). High molecular mass standards were run in the same gels. To assay denaturant conditions, 4–20% SDS–PAGE containing 0.1% SDS was carried out; samples were denatured at 100 °C for 10 min with and without reducing agent (2-ME). Gels were stained with Coomassie Brilliant Blue R-250 *Pomacea diffusa* PVF was also analysed by native PAGE and protein bands were quantified by calibrated scanning densitometry using the ImageJ software (Bourne and Bourne, 2010).

### Size and global shape

The molecular weight of PdPV1 was estimated by 4-20% Native-PAGE using high molecular weight standard and by SEC as previously described in Ituarte et al. (Ituarte *et al*., 2008), calibrated with thyroglobulin, ferritin, catalase, aldolase and ovoalbumin as MW standards. Protein global shape was determined by Small Angle X-ray scattering (SAXS). Assays were performed at the D02A-SAXS2 line, in the Laboratorio Nacional de Luz Sincrotron, Campinas (SP, Brazil). The scattering pattern was detected using a MARCCD bidimensional charge-coupled device assisted by Fit2d software (Hammersley, 2016). The experiments were performed using a wavelength of 1.448 Å. The distance between the sample and the detector was 1044 mm, allowing a Q-range between 0.012 and 0.25 Å^-1^ (Dmax =260 Å). Temperature was controlled using a circulating water bath, and kept to 25 °C. Each individual run was corrected for sample absorption, photon flux, buffer scattering, and detector homogeneity. At least three independent curves were averaged for each single experiment, and buffer blank scattering was subtracted. To rule out a concentration effect in the data, SAXS experiments with a protein concentration range of 3.0–0.2 mg/mL were performed. Data was analysed using the ATSAS package 2.6.0 (Petoukhov *et al*., 2012). The low-resolution model of PdPV1 was obtained from the algorithm built with DAMMIN and DAMMIF programs (Svergun, 1999). The average of the best 10 models fitting the experimental data was obtained with DAMMIF. Raw data was deposited in the Small Angle Scattering Biological Data Bank SASBDB.

### Absorption spectra

UV-Vis absorption spectra of PdPV1 were measured between 250-650 nm using an Agilent 8453 UV/Vis spectrophotometer at room temperature. Three spectra were recorded for each sample and its buffer contribution (50 mM NaHPO4, 150 mM NaCl pH 7.0) was subtracted.

### Carbohydrate content

Carbohydrate content of PdPV1 was determined following the phenol/sulfuric method described by (Dubois *et al*., 1956) using D-glucose as standard. Samples were diluted to a final concentration of 50 μg/mL (final volume 150 μL). All samples were treated with a 5% (v/v) phenol solution and concentrated sulfuric acid. After 30 min at 37 °C absorbance was read at 485 nm to determine hexose and pentose content.

### Determination of glycan motifs using lectin dot blot assay

A set of 13 biotinylated lectins were used, namely ConA, WGA, PSA, PNA, JAC, RCA 1, SBA, DBA, UEA, PVL, LCA, ECL and BSL 1. Lectins were reconstituted in 10 mM Buffer Hepes, 0.1 mM Cl2Ca and then diluted in 3% (w/v) BSA in PBS. Dots containing 3 μg of PdPV1 were spotted on nitrocellulose strips and incubated for 30 min at room temperature. Membranes were then blocked with 3% (w/v) BSA in PBS overnight at 4 °C and incubated for 1.30 h with 0.02 mg of the lectin. Then they were incubated with horseradish peroxidase streptavidin conjugate diluted 1:3 in water for 20 min at 25 °C. Between every step strips were washed 5 times with 0.1 % Tween in PBS for 3 min. Positive reactions were visualized by electrochemiluminiscence using the ABC system following manufacturer instructions.

### Immunoblotting

Cross reactivity of PdPV1 with anti-PcOvo and anti-PsSC polyclonal antibodies (Dreon 2003 and Ituarte 2008, respectively) was analysed by Western Blot assays (WB). Briefly, purified PdPV1 as well as PcOvo and PsSC perivitellins were run in 4-20% SDS-PAGE and then transferred onto nitrocellulose membranes using 25 mM Tris–HCl, 192 mM glycine, 20% (v/v) methanol, pH 8.3 buffer in a Mini Transblot Cell (Bio Rad Laboratories, Inc.) at 100 V for 1 h. Membranes were blocked with 5% (w/v) casein in PBS overnight at 4°C and then incubated with each polyclonal antibody at different dilutions in 3% (w/v) casein-PBS for 2 h. Finally, a goat anti-rabbit horseradish-peroxidase conjugated secondary antibody was prepared in 3% (w/v) casein-PBS, incubated for 1h with the membranes and the reaction was revelled by luminol-coumaric reaction.

### Subunit sequences and bioinformatic analysis

Internal amino acid sequences of purified PdPV1were obtained by mass spectrometry at the CEQUIBIEM (UBA–CONICET, Argentina). N-terminal sequences of subunits were determined by Edman degradation at the Laboratorio Nacional de Investigación y Servicios en Péptidos y Proteínas (LANAIS-PRO, UBA-CONICET, Argentina). These partial sequences were used to search the *P. diffusa* albumen gland transcriptome in AmpuBase (http://www.comp.hkbu.edu.hk/~db/AmpuBase/#&panel1-3) for the corresponding full protein sequences. Sequence alignment was performed using MUSCLE. PdPV1 sequences were analysed using PsiBlast, FFAS server and Hhpred. A search in PROSITE was performed to look for conserved domains. For low complexity regions, we used SEG server. The signal peptide cleavage sites in the amino acid sequences were predicted using the SignalP 4.1 server. The theoretical molecular weight and isoelectric point of each mature subunit were estimated using the ProtParam tool-Expasy server. Potential phosphorylation and glycosylation sites were predicted with DISPHOS 1.3 (Iakoucheva *et al*., 2004) and NetNGlyc 1.0 (http://www.cbs.dtu.dk/services/NetNGlyc) and NetOGlyc 4,0 (Steentoft *et al*., 2013) servers, respectively. PdPV1, PsSC, PcOvo and PmPV1 subunit sequences were subjected to phylogenetic analysis using the maximum likelihood method in MEGA X (Kumar *et al*., 2018).

### Chemical denaturation

Protein chemical denaturation by guanidine hydrochloride (GdnHCl) of PdPV1 was studied following intrinsic tryphtophan fluorescence emission. Increasing concentrations (0–6.5 M) of GdnHCl buffered with 0.2 M Tris-HCl were incubated with the samples (final concentration 0.15 g/L) at pH 7.4. Fluorescence spectra were recorded on a spectrofluorometer Fluorolog 3 Horiba (Jobin Yvon Technology, Moulton Park Northampton, UK) at 25 °C. Excitation was set at 295 nm (slit of 4 nm) and recorded from 300 to 410 nm (slit of 5 nm). Three spectra were recorded, averaged, and the corresponding buffer contribution subtracted for each sample. The protein unfolded fraction (*f*u) was calculated using the fluorescence intensity at 327.9 nm and assuming a two-state unfolding process. The equilibrium reached in each GndHCl concentration allows the calculation of an equilibrium constant K=*f*u/(1-*f*u) and the Gibb’s free energy for the unfolding reaction in terms of this mole fractions (ΔG^0^= −RT lnK) were calculated. The dependence of ΔG^0^ on GndHCl concentration can be approximated by the linear equation ΔG^0^=ΔG^0^_H2O_ – m[GndHCl], where the free energy of unfolding in the absence of denaturant (ΔG^0^_H2O_) represents the conformational stability of the protein. The GdnHCl concentration in which half of the protein is unfolded (Cm) was estimated as a function of denaturant concentration from the linear extrapolation method.

### Resistance to sodium dodecyl sulphate

Denaturation induced by SDS is used to identify proteins whose native conformations are kinetically trapped in a specific conformation because of an unusually high unfolding barrier that results in very slow unfolding rates (Jayaprakash and Surolia, 2017). To test PdPV1 resistance to denaturation by SDS, an experiment following the Manning and Colón procedure was performed (Manning and Colón, 2004). In short, the protein in Laemmli sample buffer (pH 6.8) containing 1% SDS was either boiled for 10 min or unheated. Then samples were analysed by 4–20% SDS gel electrophoresis and visualized with Coomassie Brillant Blue.

### Structural stability against temperature and pH

Three hundred μL of PdPV1 (0.28 g/L) in 50 mM Na_2_HP0_4_, 150 mM NaCl buffer pH 7.0 were used for the assays. For thermal stability assays the sample was subjected to rising temperatures from 25 °C to 85 °C and, to assess its stability at different pH values, the protein was incubated in buffers ranging from pH 2.0 to 12.0 overnight at 4 °C. Buffer formulations were taken from Merril (1990). PdPV1 tertiary structure was monitored by absorbance and fluorescence spectroscopy and global shape and size by SAXS. Absorbance was measured between 250-800 nm with an Agilent 8453 spectrophotometer and the 4^th^ derivative spectra calculated. Florescence emission was recorded from 300-420 nm in a Varian Cary Eclipse spectrofluorometer (Varian Inc., Australia), excitation set to 295 nm with a slit of 8 nm, in 5 mm optical path quartz cells. For both techniques three spectra were recorded and averaged for each sample and its corresponding buffer blank was subtracted. SAXS experiments were performed as described above, except for thermal stability of PdPV1 for which the sample holder temperature was controlled using a water circulating bath between 25 to 85 °C. The effect of extreme thermal conditions was analysed by boiling PdPV1 for 15 min and evaluating the oligomer integrity using native (non-denaturing) gel electrophoresis and the carotenoprotein fine spectra using absorbance spectrophotometry.

### Capacity to withstand in vivo and in vitro gastrointestinal digestion

PdPV1 resistance to protease degradation was studied *in vitro* and *in vivo* as previously described (Dreon, Ituarte and Heras, 2010; Pasquevich *et al*., 2017). For the simulated gastrointestinal digestion, the purified protein (105 μg) in phosphate buffer was first subjected to pepsin at a ratio enzyme:substrate 1:20 (w/w), at 37 °C in a simulated gastric fluid (SGF) which contains 150 mM NaCl, pH 2.5. Serine bovine albumin (BSA) was used as control. Aliquots of the sample and control were taken after 60 min and 120 min of incubation. To stop the reaction, pH was raised to 8.5 with 150mM Tris-Cl buffer and samples were kept a −20 °C until processing. The remnant of the solutions was then incubated with trypsin to simulate the gut phase in a ratio enzyme:substrate 1:2.7 (w/w) at 37 °C in SFG. The pH of the solution was adjusted with 0.1 M NaOH, and 0.25 M Taurocolic acid and Tris-HCl 0.15 M buffer, pH 8.5 were added. Aliquots of the sample and control were taken at the same times as in gastric phase and stored at −20 °C. Finally, all samples were boiled at 100 °C for 10 min and run in 4-20 % SDS-PAGE gradient to check the integrity of the protein.

To evaluate PdPV1 resistance to *in vivo* gastrointestinal digestion, animals were orally administrated with 0.6 mg of PdPV1 in 50 mM Na2HPO4, 150 mM NaCl buffer and their faeces were collected every hour during 4 h after oral administration. During and before the experiment animals were fed *ad libitum*. Control BALB/c mice were only given buffer. Faecal extracts were resuspended in EDTA 30 mM pH 8.4 in PBS with protease inhibitor 1:100 (Sigma) and homogenized in a Teflon homogenizer before centrifugation at 13,000 g at 4 °C for 10 min. Samples were run in 4-20 % Native-PAGE gradient and stained with Coomassie Brillant Blue to check for the presence of the protein in the faeces. WB was performed as stated in immunoblotting section to visualize the protein subunits using purified PdPV1 as control.

### Neurotoxicity

To evaluate whether *P. diffusa* eggs contain neurotoxins as reported in other *Pomacea* species (Heras *et al*., 2008; Giglio *et al*., 2016), groups of 5 mice were intraperitoneally injected with either 200 μL of PBS or 4.6 mg/kg (200 μL) of PVF. Mice behaviour was observed along 96 h and behavioural changes or neurological signs recorded. Special attention was given to those changes already reported for *Pomacea* neurotoxins including weakness and lethargy, tachypnea, hirsute hair, abduction of the rear limbs preventing them from supporting their own weight and spastic movements of tail muscles.

### Glycan binding specificity using Glycan array

Glycan binding specificity of PdPV1 was determined at the Core H of the Consortium for Functional Glycomics (http://www.functionalglycomics.org Emory University, Atlanta, GA, USA). The protein was fluorescently labelled using the Alexa Fluor 488 Protein Labelling kit according to the manufacturer’s instructions. Labelled PdPV1 was assayed on a glycan array which comprised 585 glycan targets (version 5.4) and the data analyzed as described in Ituarte et al (Ituarte *et al*., 2018).

### Hemagglutination assays

PdPV1 lectin activity was tested by hemagglutination of red blood cells (RBC) from rabbit, human, goat, horse, rat and chicken. Cells were obtained from the animal facilities of National University of La Plata (UNLP). Blood was taken by venous puncture and collected in sodic EDTA 300 mM in sterile conditions. Human RBC of type A, B and 0 from healthy donors were donated by the Institute of Hematology (Ministerio de Salud, Provincia de Buenos Aires, Argentina). Erythrocytes were prepared as stated in Ituarte et al. (Ituarte *et al*., 2012). 50 μL of two-fold serial dilutions of PdPV1 in PBS were incubated with an equal volume of 2% (v/v) erythrocytes in PBS in U-shaped microtiter plates (Greiner Bio One, Germany) at 37 °C for 2 h. The initial protein concentration was of 0.4 mg/mL.

PdPV1 carbohydrate affinity was inferred by comparing the inhibitory activity of the addition of fucose, N-acetyl-glucosamine, N-acetyl-galactosamine, glucosamine, galactosamine, mannose, galactose and glucose on hemagglutination. Two-fold serial dilutions of purified protein starting from 0.4 mg/mL were made in 200 mM of each carbohydrate and incubated for 1 h at 37 °C. Later, 50 μL of 2 % rabbit erythrocytes suspension in PBS were added to these solutions in a microtiter plate U shaped and after 2 h hemagglutination activity was verified. The inhibitory capacity was expressed as the percent of inhibition, compared with the agglutination titter of PdPV1 without inhibitors: Inh% = [100 - (Ti 100)/T], where T represents the hemagglutinating titter of the lectin without inhibitors and Ti titter with inhibitors. In some experiments, RBC were pre-treated with trypsin 0.1 mg/ml or neuraminidase 0.1 U/ml (BioLabs, New England) before agglutinating analysis.

#### Statistical analysis

Data were analyzed by one-way analysis of variance (ANOVA). When p values were < 0.05, the significance between groups was estimated by the Tukey’s test.

## Results

### Structure

PdPV1 was purified and isolated by ultracentrifugation and HPLC. After NaBr gradient ultracentrifugation of the egg soluble fraction, two protein fractions were obtained: a colourless fraction of ~1.22 g/mL and a coloured fraction of ~1.24 g/mL (Fig 1A). When the coloured fraction was subjected to SEC, two peaks were observed (Fig 1B). The larger one was a carotenoprotein we named PdPV1, while the other is a PdPV1 aggregate, having both peaks the same apoprotein composition in SDS PAGE (Fig 1B inset). Native (non-denaturing) PAGE of PVF proteins of *P. diffusa* eggs (Fig 1C) showed that PdPV1 is the most abundant representing ca. 60% of the total protein. The estimated molecular mass by SEC was 256 kDa, although the apparent mass by non-denaturing PAGE was ~470 kDa (Fig 1C). PdPV1 is composed of several subunits ranging from 23-33 kDa as determined by SDS PAGE under reducing conditions (Fig 1B). These subunits are not held together by disulphide bonds as SDS-PAGE performed under non-reducing conditions indicate (not shown). The absorption spectrum of PdPV1 has the carotenoid-characteristic bands, with a single absorption maximum at 382 nm (Fig 1D). However, unlike the other apple snail carotenoproteins, its intensity (and hence the egg coloration) is very weak. Total carbohydrate content of PdPV1 represents 8.9 % (by wt), which probably resulted in an inaccurate molecular weight estimation by native PAGE. The protein glycosylation pattern was determined by lectin dot-blot (Table 1). The absence of reactivity with PSA together with positive reactivity of ConA suggest the presence of trimannoside core glycans, while DBA, SBA and PNA positive reactivity indicates the presence of galactosides. PdPV1 has non-substituted T-antigen as suggested by PNA positive reactivity. The global shape of the particle in solution was determined by SAXS. A gyration radius (Rg) of 45.5 Å was estimated from the Guinier plot. PdPV1 is a globular protein as evidenced from the bell-shaped form of the curve in the Kratky plot (see SASBDB). A low-resolution 3D-model for PdPV1 is depicted in Fig 1E. SAXS raw data were deposited in SASBDB repository (https://www.sasbdb.org/data Accession Code SASDG75).

**Figure 1:**
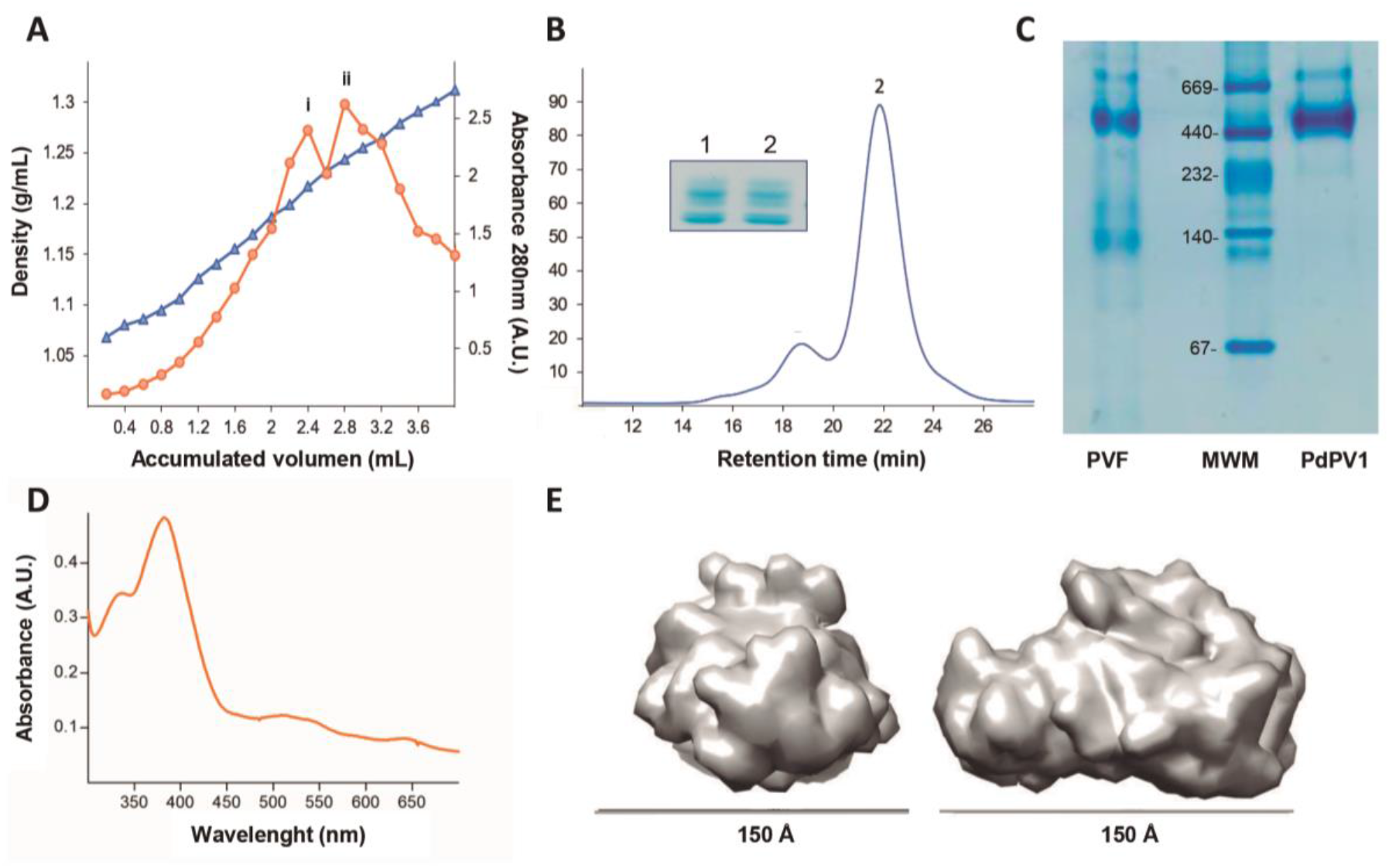
Purification and structure of PdPV1. A) Protein (orange circles) and hydration density (blue triangles) profile of *P. diffusa* egg soluble fraction; (i) uncoloured and (ii) coloured fraction. B) Size exclusión chromatogram of ii coloured fraction, (1) Agregate, (2) PdPV1. Inset: SDS-PAGE of 1 and 2 peaks showing the same apoprotein composition. C) Native PAGE of the egg soluble fraction (PVF) and purified PdPV1. MWM: Molecular weight markers. D) Absorption spectrum of purified PdPV1. E) *Ab-initio* model of SAXS data of PdPV1 by DAMMIF (representative of best model cluster).

**Table 1.** Acronyms and specificity of lectins employed in PdPV1 oligosaccharides recognition

| Acronym | Binding specificity | PdPV1 |
| --- | --- | --- |
| ConA | Man α3[Man α6] man; terminal αMan; oligomannose and hybrid type N-glycans | + |
| WGA | Neu5Acα3/6/8 R | + |
| PSA | N-acetylchitobiose-linked Fuc | - |
| PNA | Terminal Gal β13-GalNAc | + |
| JAC | GalNAc | + |
| RCA-1 | Terminal βGal | + |
| SBA | Tn antigen; terminal $\alpha$ GalNAc | + |
| DBA | GalNAc α1,3 GalNAc/Gal | + |
| UEA | $\alpha$(1,2) L-Fuc | + |
| PVL | GlcNAc Neu5Ac α26-Gal | - |
| LCA | Bi- and triantennary complex-type N-glycan | + |
| ECL | Terminal LacNAc | - |
| BSL-1 | αGalNAc; αGal | + |

Major specificities (Goldshtein and Hayes, 1978). Fuc, L-Fucose; Gal, galactose; GalNAc, N- acetyl galactosamine; Glc, glucose; GlcNAc, N-acetyl glucosamine; Man, Mannose; NeuNAc, acetyl neuraminic acid (sialic acid).

The internal sequences, together with three N-terminal aminoacid sequences (Table S1) were used to identify six full length subunits (Fig. S1A) in the *P. diffusa* albumen gland transcriptome, hereafter referred as PdPV1-1 to 6. All six deduced aminoacid sequences have a signal peptide and a theoretical molecular mass around 21 kDa in agreement with the MW estimation by SDS-PAGE. All subunits have at least one potential phosphorylation site as follows: PdPV1-1 has one Ser and two Tyr; PdPV1-2 has 50% of Tyr predicted to be phosphorylated; PdPV1-3 has two Thr and one Tyr; PdPV1-2 six predicted sites (1 Ser, 2 Thr, 3 Tyr); PdPV1-5 two Ser; PdPV1-6 one Ser and one Tyr. Regarding potential glycosylation sites on PdPV1, while the 6 subunits present at least one N-glycosylation site, predicted O-glycosylation sites are only present in PdPV1-4, PdPV1-5 and PdPV1-6 (Figure S1A). SEG server indicates the absence of low-complexity regions in these sequences. Sequence alignment of the six PdPV1 subunits is shown in Fig. S1B.

An NCBI nr database search revealed no homology with any known sequences other than those of *Pomacea* perivitellins. Using PsiBLAST search revealed homologues only within the genus; no conserved domains were identified by PROSITE. In addition, no sequence similarity with any known lectin could be found for any of the six PdPV1 subunits, except for PsSC lectin from the sister species *P. scalaris*.

### Evolutionary relationships among Pomacea perivitellins

A phylogenetic analysis performed including all *Pomacea* perivitellin subunits sequences (Fig 2A) indicates that PdPV1 subunits did not group together but each one grouped with those of the other species into six separate clusters. Each orthologous sequence cluster among PdPV1, PsSC, PcOvo and PmPV1 was separated into two subgroups that corresponded to *bridgesii* and *canaliculata* clades (Fig. 2B). Comparison of amino acid sequences within each orthologue cluster indicate high similarity. However, among the 6 PdPV1 subunits, the similarity between any two subunits was significantly lower. The immune cross reactivity among PdPV1, PcOvo and PsSC was evaluated by Western blot assays. Anti-PsSC polyclonal antibodies strongly recognized each of the six PdPV1 subunits, whereas anti-PcOvo polyclonal antibodies weakly cross-reacted with only a few PdPV1subunits (Fig 2C).

**Figure 2:**
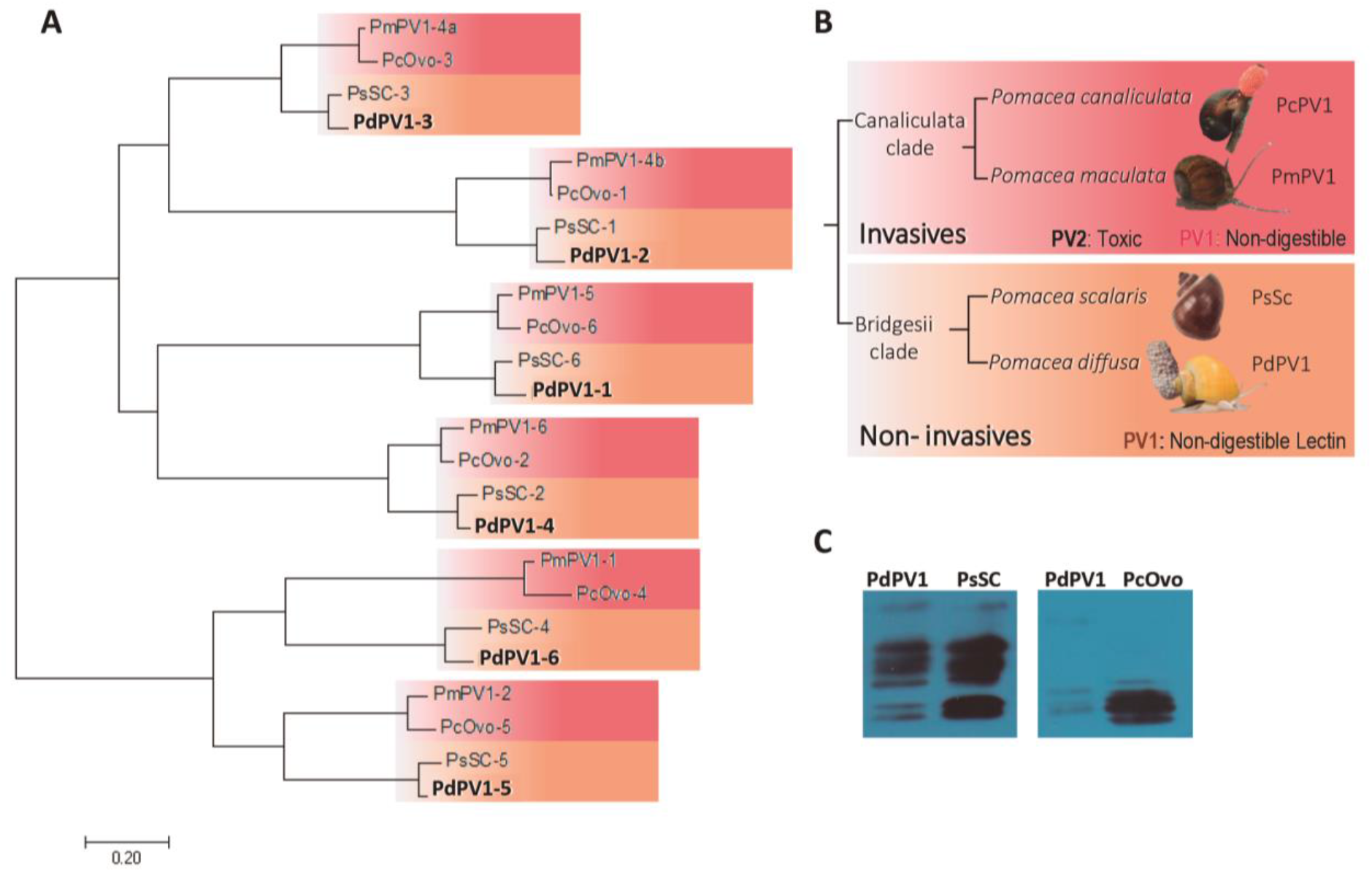
Phylogenies of PV1perivitellins subunits versus morphological phylogenies of *Pomacea* Canaliculata and Bridgesii clades. A) Maximum-likelihood phylogram obtained with RAxML under the best partition model for Pomacea ^‥^PV1^‥^ perivitellins. Values at nodes represent maximum-likelihood bootstrap percentages under the best partition model using RAxML (BPRAxML) and IQ-TREE (BPIQ-TREE) and clade posterior probabilities under the best partition model using MrBayes (PPMrBayes) and the CAT-GTR mixture model using PhyloBayes (PPPhyloBayes). In bold proteins from this study. (B) Phylogeny of *Pomacea canaliculata* and *bridgesii* clades (after Hayes, 2009) indicating species, invasiveness and some of the perivitelline defenses present in each. (C) Cross reactivity of PdPV1 with polyclonal antibodies anti-PsSC (left panel) and anti-PcOvo (right panel).

### Lectin activity

PdPV1 showed hemagglutination activity against rabbit, rat and human red blood cells (RBC) and non-detectable activity toward RBC from other species. This activity was largely inhibited by N-Acetyl-galactosamine, galactosamine and galactose, while preincubation of rabbit RBC with neuraminidase did not block hemagglutination. Glycan specificity and relative affinity was further evaluated by a glycan array screening of PdPV1. Out of 523 unique glycan structures, we observed significant binding of PdPV1 to a range of structures that included 2,8-linked N-acetylneuraminic acid, a common sialic acid in vertebrate ganglioside motifs. In fact, PdPV1 recognizes with high affinity GD1b, GT2, GD2, GT1b, GD1a gangliosides. Interestingly, PdPV1 higher affinity scores were against sialylated structures that featured a GalNAcb1-4Gal motif. PdPV1 affinity for these GalNAc-Gal would explain the presence of aggregated PdPV1 lectin observed in the chromatographic analysis (Fig.1B) because these glycans, which predominate in the glycan decoration of PdPV1 (Table 1) would promote self-aggregation. Very few examples of PdPV1 binding to complex N-linked glycans were observed. Glycans with higher scores are listed in Table 2.

**Table 2.**
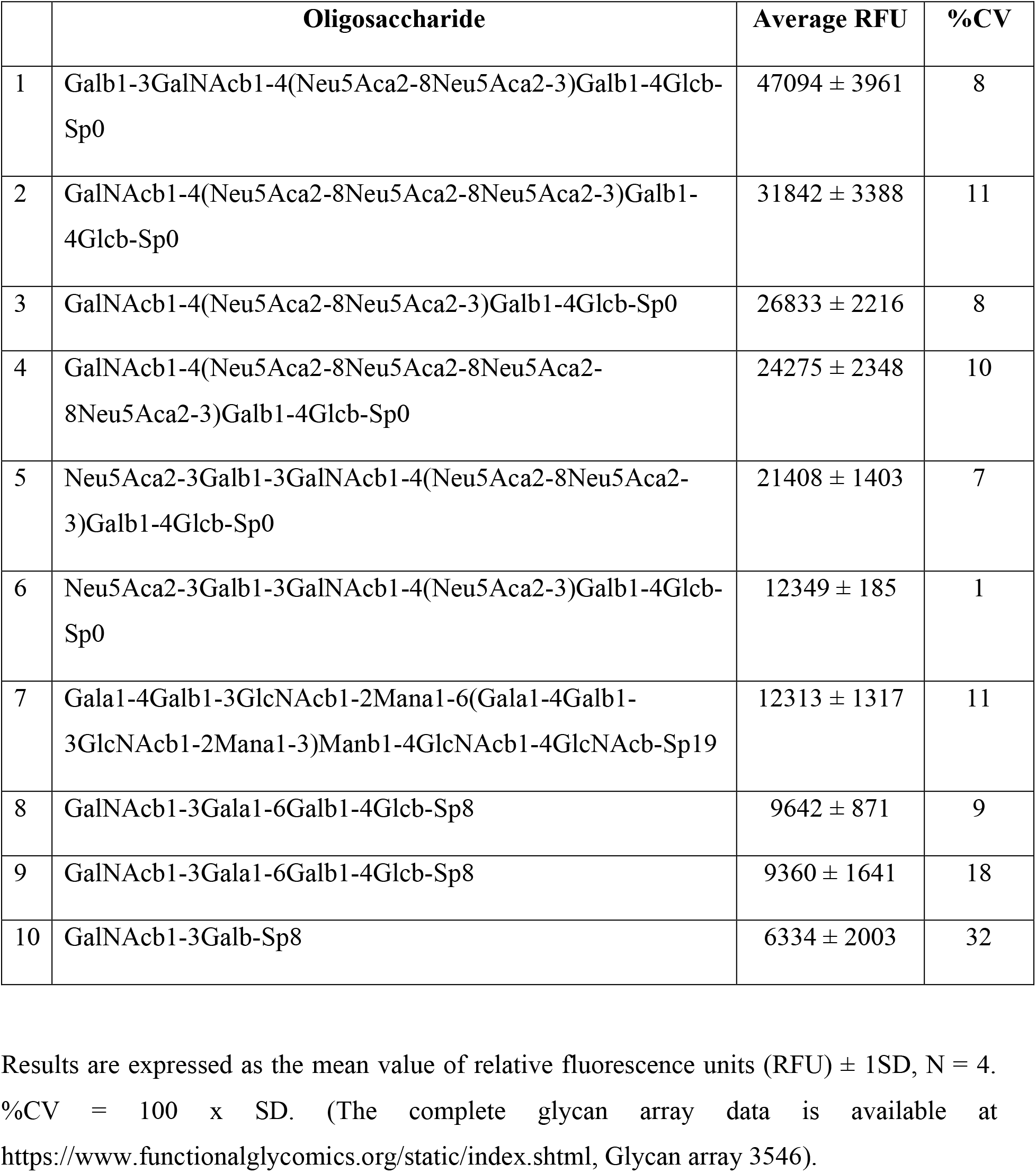
Main glycan structures recognized by PdPV1

|  | **Oligosaccharide** | **Average RFU** | **%CV** |
| --- | --- | --- | --- |
| 1 | Galb1-3GalNAcb1-4(Neu5Aca2-8Neu5Aca2-3)Galb1-4Glc-Sp0 | 47094 ± 3961 | 8 |
| 2 | GalNAcb1-4(Neu5Aca2-8Neu5Aca2-8Neu5Aca2-3)Galb1-4Glc-Sp0 | 31842 ± 3388 | 11 |
| 3 | GalNAcb1-4(Neu5Aca2-8Neu5Aca2-3)Galb1-4Glc-Sp0 | 26833 ± 2216 | 8 |
| 4 | GalNAcb1-4(Neu5Aca2-8Neu5Aca2-8Neu5Aca2-8Neu5Aca2-3)Galb1-4Glc-Sp0 | 24275 ± 2348 | 10 |
| 5 | Neu5Aca2-3Galb1-3GalNAcb1-4(Neu5Aca2-8Neu5Aca2-3)Galb1-4Glc-Sp0 | 21408 ± 1403 | 7 |
| 6 | Neu5Aca2-3Galb1-3GalNAcb1-4(Neu5Aca2-3)Galb1-4Glc-Sp0 | 12349 ± 185 | 1 |
| 7 | Gala1-4Galb1-3GlcNAcb1-2Mana1-6(Gala1-4Galb1-3GlcNAcb1-2Mana1-3)Manb1-4GlcNAcb1-4GlcNAcb-Sp19 | 12313 ± 1317 | 11 |
| 8 | GalNAcb1-3Gala1-6Galb1-4Glc-Sp8 | 9642 ± 871 | 9 |
| 9 | GalNAcb1-3Gala1-6Galb1-4Glc-Sp8 | 9360 ± 1641 | 18 |
| 10 | GalNAcb1-3Galb-Sp8 | 6334 ± 2003 | 32 |

Results are expressed as the mean value of relative fluorescence units (RFU) ± 1SD, N = 4.
%CV = 100 x SD. (The complete glycan array data is available at
https://www.functionalglycomics.org/static/index.shtml, Glycan array 3546).

### Neurotoxicity

Intraperitoneal injection of PVF had no effect on mice survival. Moreover, after 96 h mice did not show neither neurological signs nor behavioural changes.

### Structural stability

Structural stability of PdPV1 at different pH values was monitored by fluorescence and absorption spectroscopy and by SAXS. Absorption and fluorescence spectroscopy showed that only at pH 2.0 the protein partially alters its conformation as seen by an intensity decrease in the carotenoid absorbing (Fig 3A) and a blue shift in the 4^th^ derivate of the spectra (Fig 3B). No changes in tryptophan fluorescence emission spectra were recorded (Fig 3C), indicating that the protein remains properly folded up to pH 2.0, where a slight red shift of the emission maximum was observed. However, changes at pH 2.0 might be consider small changes in protein structure because SAXS experiments showed that the particle shape remains globular in the pH 2.0-12.0 range (Fig 3D) with no changes of its Rg except for a small reduction at pH 12.0 (Fig 3E). Regarding thermal stability, no structural perturbations in the conditions assayed could be detected in PdPV1 neither by absorption (Fig 4A and B) nor fluorescence spectroscopy (Fig 4C), indicating protein stability up to 85 °C. These results are supported by SAXS although at 85°C a partial loss of protein globularity occurs (Fig 4D) associated with a slight Rg increase (Fig 4E). Moreover, extreme thermal treatment of PdPV1 (boiled for 15 min) did not affect its oligomer integrity as seen by native PAGE (Fig 4F inset).

**Figure 3:**
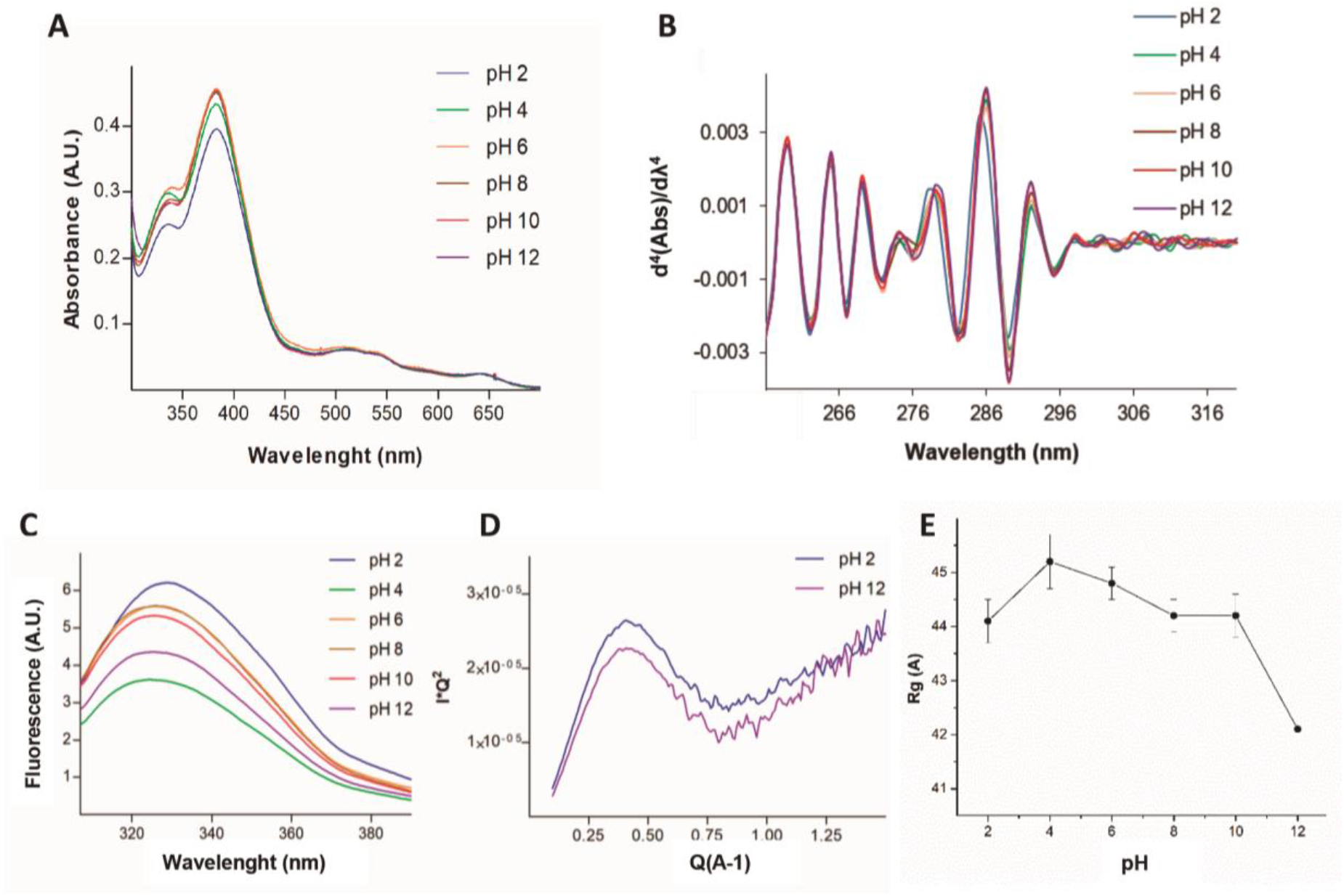
Effect of pH on PdPV1 structural stability. A) Absorption spectra. B) Fourth derivative absorption spectra. C) Intrinsic fluorescence emission spectra and D) Kratky plots of PdPV1 at pH 2.0 (blue) and 12.0 (violet). E) Gyration radii at different pH values.

**Figure 4:**
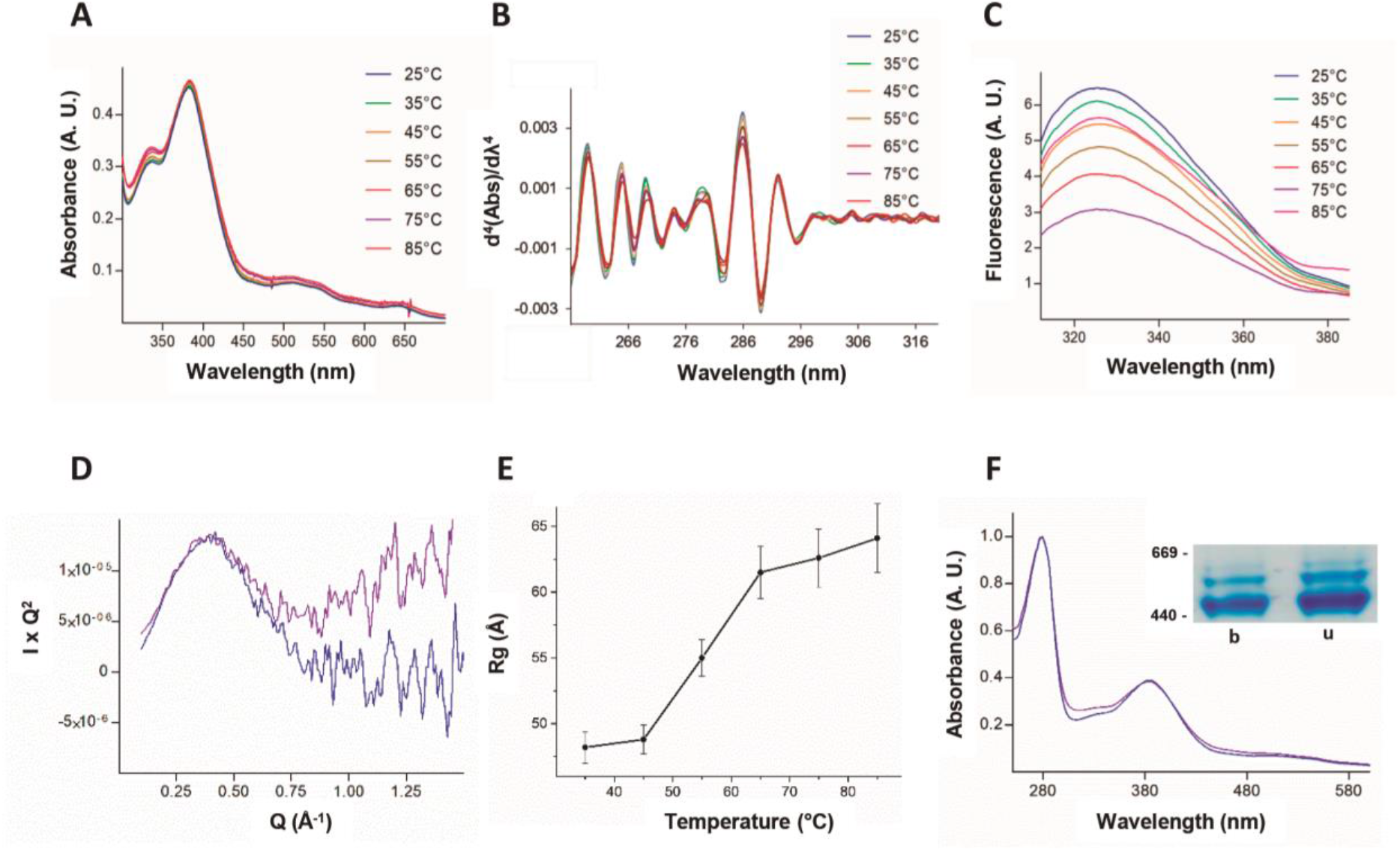
Effect of temperature on PdPV1 structural stability. A) Absorption spectra. B) Fourth derivative absorption spectra. C) Intrinsic fluorescence emission spectra. D) Kratky plots obtained from SAXS data at 25 °C (blue) and 85 °C (violet). E) Gyration radii at different temperatures. F) Absorption spectra of unboiled (blue) and boiled (violet) PdPV1, inset: Native 4-20% PAGE. Lane b: PdPV1 boiled for 15 min; lane u: PdPV1 unboiled.

Chemical stability of PdPV1 was studied by equilibrium unfolding experiments. The protein unfolded fraction vs [GndHCl] is depicted in figure 5A, showing cooperative unfolding transitions. The unfolding process fits to a two-state model with the protein completely unfolded from 4.5 M GnHCl onwards. The calculated standard free energy for the unfolding process was 1.14 Kcal.mol^-1^, and the *m* value, which characterized the influence of the chaotrope on the process, was 338.7 cal.mol^-1^.M^-1^. Finally, the midpoint unfolding GndHCl concentration for PdPV1 was 3.4 M. Kinetically stable proteins show no alteration on their electrophoretic mobility when incubated with SDS; they need to be heated to be disassembled into their subunits. In this regard, PdPV1 did not changed it migration pattern after incubation with SDS. Only when PdPV1 was boiled in the presence of SDS the oligomer disassembled into its 6 subunits as evidenced by SDS PAGE (Fig 5B).

**Figure 5:**
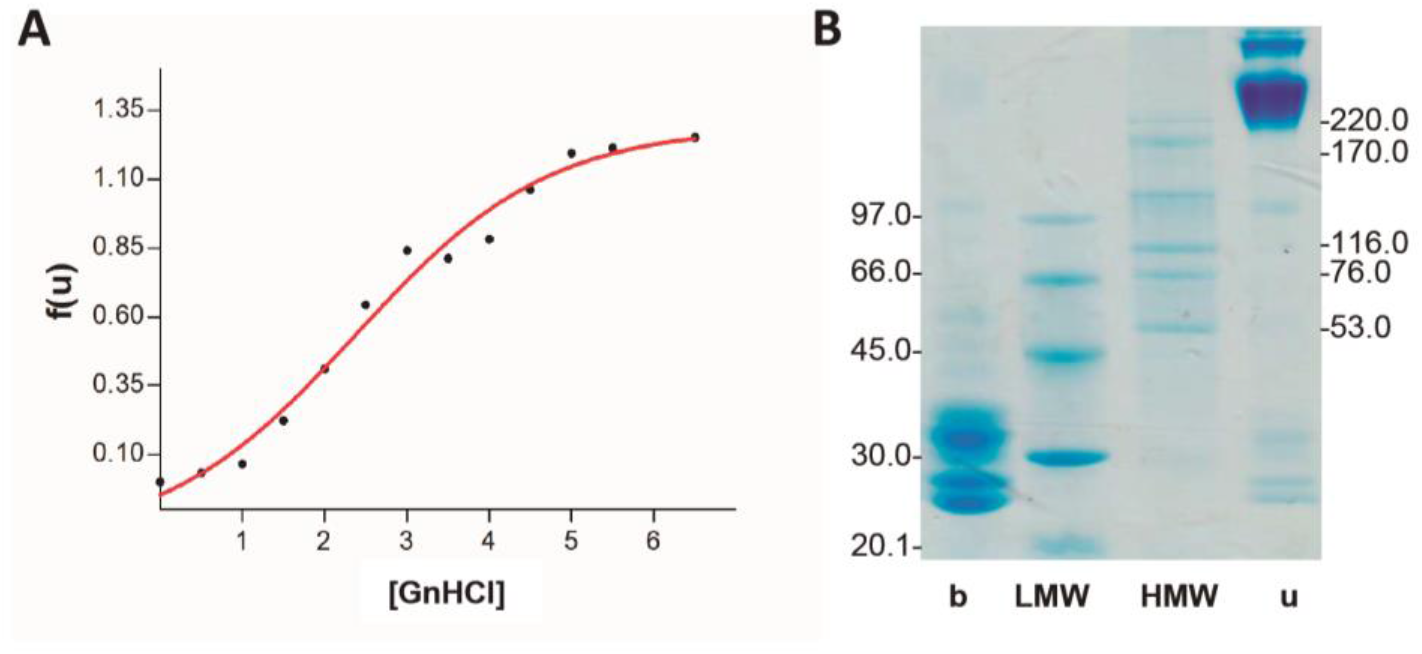
Structural stability of PdPV1. A) Chemical stability evaluated by the unfolding induced by GnHCl. B) Kinetic stability evaluated with an SDS-resistance assay. The same PdPV1 sample was unheated (u) or boiled (b) in the presence of SDS for 10 min immediately prior to be loaded into the gel. LMW and HMW: low and high molecular weight markers, respectively.

The particle was also very resistant to enzymatic digestion, withstanding a simulated gastrointestinal digestion for 2 h. The integrity of the protein after gastric and duodenal phases was analysed by SDS-PAGE showing no significant alterations in its electrophoretic migration (Fig 6A). Furthermore, when the protein was administered to mice, the protein was recovered almost intact in the faeces as shown by Native PAGE and WB analysis indicating that PdPV1 can pass through the digestive system without altering its structure (Fig 6B and C).

**Figure 6:**
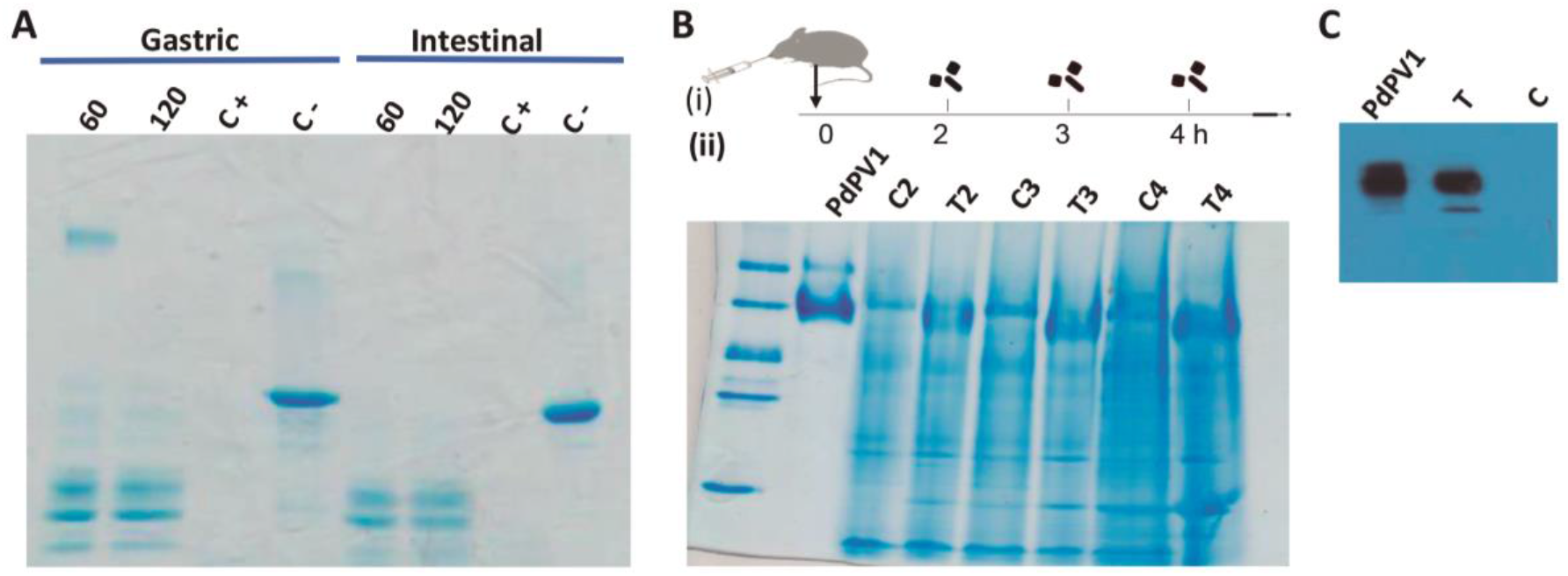
*In vitro* and *in vivo* digestibility of PdPV1. A) Simulated gastrointestinal digestion analyzed by SDS-PAGE. Gastric phase: PdPV1 after incubation with pepsin in SGF for 60 and 120 min (60 and 120): C+, C-: positive and negative controls of BSA digestion with and without pepsin, respectively. Intestinal phase. PdPV1 incubated with trypsin for 60 (60) and 120 (120) min; C+ and C-positive and negative controls of BSA digestion with (C+) and without (C-) addition of trypsin. B) *In vivo* digestibility of PmPV1. (i) Diagram of experimental protocol showing oral administration of PmPV1 and faeces collection times. (ii) Analysis of faecal proteins by Native PAGE. MW: Molecular weight markers; PdPV1: purified PdPV1; C2, C3, C4, and T2, T3 and, T4: control and faeces collected after 2,3 and 4 h, respectively. C) Immune-detection by Western blot of PdPV1 in faeces collected after 4h mice were gavaged with buffer (C) or with 0.6 mg PdPV1 (T).

## Discussion

During *Ampullariidae* transition from water to land*, Pomacea* eggs faced the harsh environmental conditions of the terrestrial environment, acquiring novel defences, including highly stable, non-digestible oligomeric perivitellins that became the most abundant proteins in the PVF (Dreon, Heras and Pollero, 2003; Ituarte *et al*., 2008, 2018; Dreon, Ituarte and Heras, 2010; Pasquevich, Dreon and Heras, 2014; Pasquevich *et al*., 2017; Ip *et al*., 2019, 2020). In this study we isolated and characterized the main perivitellin of *P. diffusa* eggs, which unveiled some evolutionary trends within the genus. We found that, as in other *Pomacea*, PdPV1 is also a highly stable carotenoprotein that remains completely folded even at high temperatures, extreme pH values and high concentrations of chaotropes. Furthermore, this remarkable structural stability allows PdPV1 to be undigestible, withstanding the passage through the gastrointestinal tract if ingested by mice, indicating it could reach the digestive epithelium in a fully active conformation.

PdPV1 is composed by six polypeptide chains similar to its orthologs PcOvo, PsSC and PmPV1 (Pasquevich *et al*., 2017; Ituarte *et al*., 2018, 2019) with only moderate sequence similarities among them. It has been suggested that they were generated by an ancient gene duplication (Sun *et al*., 2012). Phylogenetic analysis of the four perivitellins, shows each PdPV1 subunit grouped separately in one of six different clades, each containing the corresponding orthologous subunit of each species perivitellin. In addition, each orthologous group presents low sequence divergence, reinforcing the suggestions by Ituarte et al. (Ituarte *et al*., 2018), that after gene duplication the advantage of having multiple copies of this gene may enhance the perivitelline synthesis during egg production, thus preventing additional gene variations. Moreover, the high expression level of these genes, almost exclusively restricted to the albumen gland of *P. diffusa* (Ip et al., 2020), further supports this hypothesis.The perivitellin family tree agreed with those based on morphology and MT COI genetic analyses (Hayes *et al*., 2015), as the subunits belonging to PsSC and PdPV1 (*bridgesii* clade) and PcOvo and PmPV1 (*canaliculata* clade) grouped separately. This was further evidenced using antibodies: while *bridgesii* clade perivitellins are recognized by an anti-PsSC polyclonal antibody, anti-PcOvo antibodies reacted weakly. These results provide the first evidence of structural differences between perivitellins from *bridgesii* and *canaliculata* clades, probably associated but not restricted to their different glycosylation patterns. In this regard, PdPV1 glycosylation pattern is quite similar to PsSC (Ituarte *et al*., 2010) but differs from those of perivitellins from the *canaliculata* clade.

For evolutionary trends to be distinguished, enough species within a group must have been studied. Thus, the characterization of PdPV1 disclose characteristics common to all apple snail carotenoproteins. They are all high molecular weight homologous oligomers formed by 6 subunits of 25-35 kDa each, with similar amino acid sequences. As antipredator molecules, they can be regarded as antinutritive proteins that lower the nutritional value of eggs as kinetically-stable and non-digestible storage proteins that survive the passage through the gut of predators unaffected (Pasquevich *et al*., 2017). However, some clade-specific differences also arise: the *bridgesii* clade with noninvasive species (*P. scalaris* and *P. diffusa*) features carotenoproteins with lectin activity that are absent in the ortholog perivitellins of the *canaliculata* clade that include the invasive species *P. canaliculata* and *P. maculata*, highlighting a clade-related association between invasiveness and type of egg defense. Assuming that PdPV1 and PsSC have similar roles, we can say they have a dual defensive function: as lectin altering predator gut morpho-physiology and as non-digestible protein lowering the nutritional value of eggs. In this regard, both PdPV1 and PsSC lectin activity (Ituarte *et al*., 2018) seems directed toward ganglioside-containing structures. The similarities between PdPV1 and PsSC in their lectin activity and primary structure, together with the lack of homology with any other lectin, led us to speculate that these perivitellins have a novel carbohydrate recognition module and could probably be a novel lectin family, although this needs further structural analysis. Lectins are essential humoral effectors in the innate immune system and play a defensive role against microorganism (Malham *et al*., 2002), a possible role for these lectins as previously suggested for PsSC (Ituarte *et al*., 2012). After ingestion, PsSC binds to enterocytes, inducing a modification of its glycosylation pattern and eventually altering the normal morpho-physiology of the rodent digestive tract, thus having adverse nutritional consequences in potential egg consumers (Ituarte *et al*., 2018). Considering that the action of lectins is predominantly triggered after recognizing their specific glycan ligands (Hirabayashi, 2014) and that both PsSC and PdPV1 bind to similar gangliosides, it is expected that binding of *Pomacea* PV1 lectins to cell surface glycolipids may be the first step of their noxious action on gut, a well-known defensive strategy in plants (Peumans and Van Damme, 1995). To the best of our knowledge, the massive accumulation in eggs of an hyperstable, non-digestible lectin, combined with protease inhibitors and its ability to adversely affect gut morphophysiology is unique among animal defensive systems against predation.

The high structural stability of PdPV1 together with its specificity for gangliosides usually overexpressed in certain types of tumors, opens a new research avenue to study PdPV1 potential biomedical applications as a tumor marker.

## Conclusions

This study contributes to our knowledge on the reproductive biology of snails. Particularly, it revealed clade-specific defensive strategies in the poisonous *Pomacea* eggs, some with carotenoproteins with lectin activity and some other with neurotoxins as the major effectors. The occurrence of different defensive strategies is related to the species’ phylogenetical position and to their invasiveness. By contrast the presence of highly stable, non-digestible storage carotenoproteins massively accumulated in the egg emerges as a common character of the whole genus *Pomacea*. Another common feature is that PV1s provide eggs with coloration, a presumed aposematic signal to deter predators from ingesting their noxious eggs. More comparative work on snail perivitellins from these and other clades is still needed to fully appreciate the unique defenses of ampulllariids and their evolutionary trends.

## Acknowledgements

MSD is member of Carrera del Investigador Científico CICBA, Argentina. HH, is member of Carrera del Investigador Científico CONICET, Argentina. TRB is a Ph.D. fellow from CONICET, Argentina. We wish to acknowledge the Protein-glycan Interaction Resource of the CFG (supporting grant R24 GM098791) and the National Center for Functional Glycomics (NCFG) at Beth Israel Deaconess Medical Center, Harvard Medical School (supporting grant P41 GM103694). We thank LNLS - Brazilian Synchrotron Light Laboratory for access to their facilities and partial financial support (Projects SAXS1-17746 and SAXS2-18846).

---

**Table S1:**
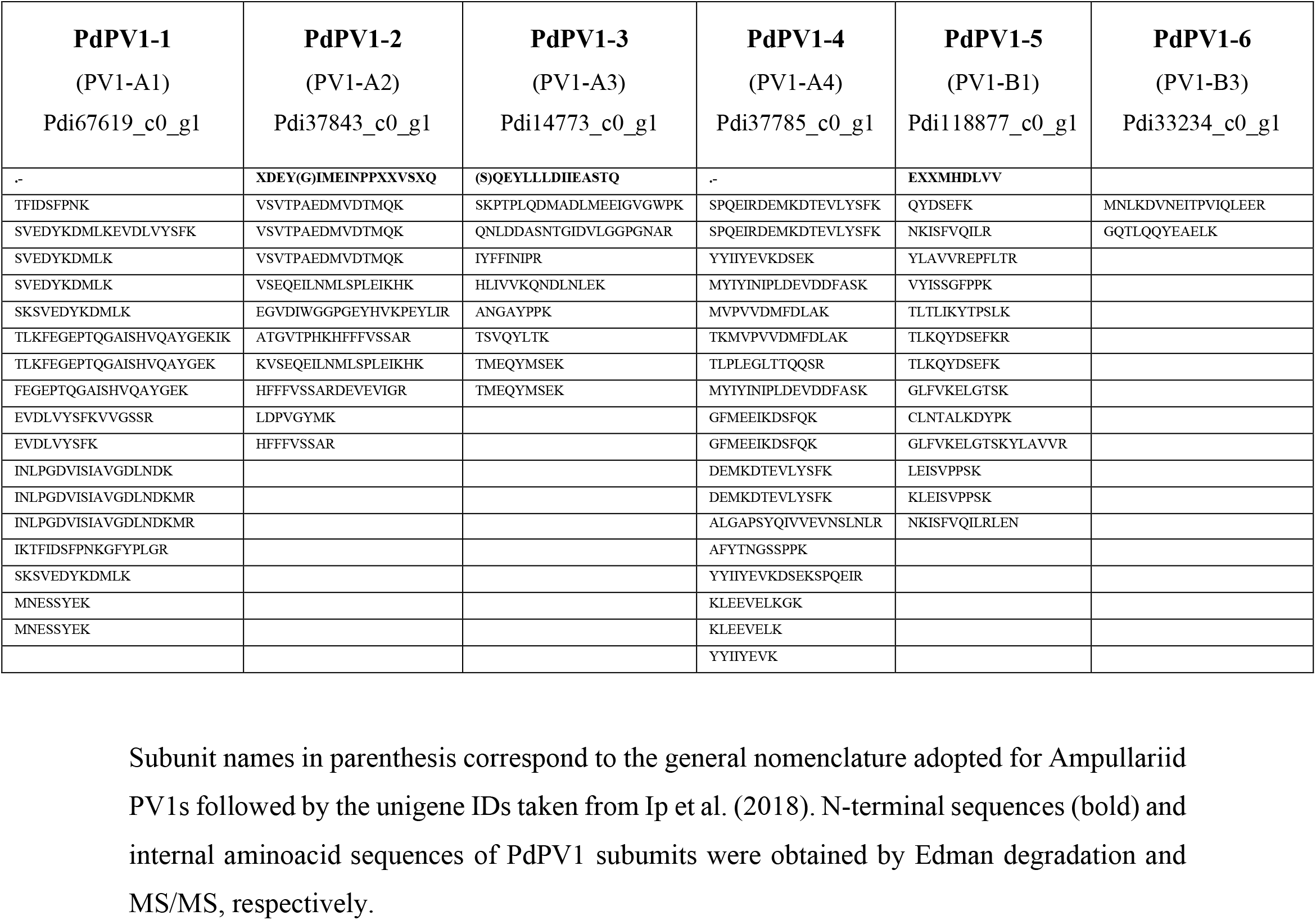
Peptide sequences employed to identify PdPV1 subunits.

**Figure S1:**
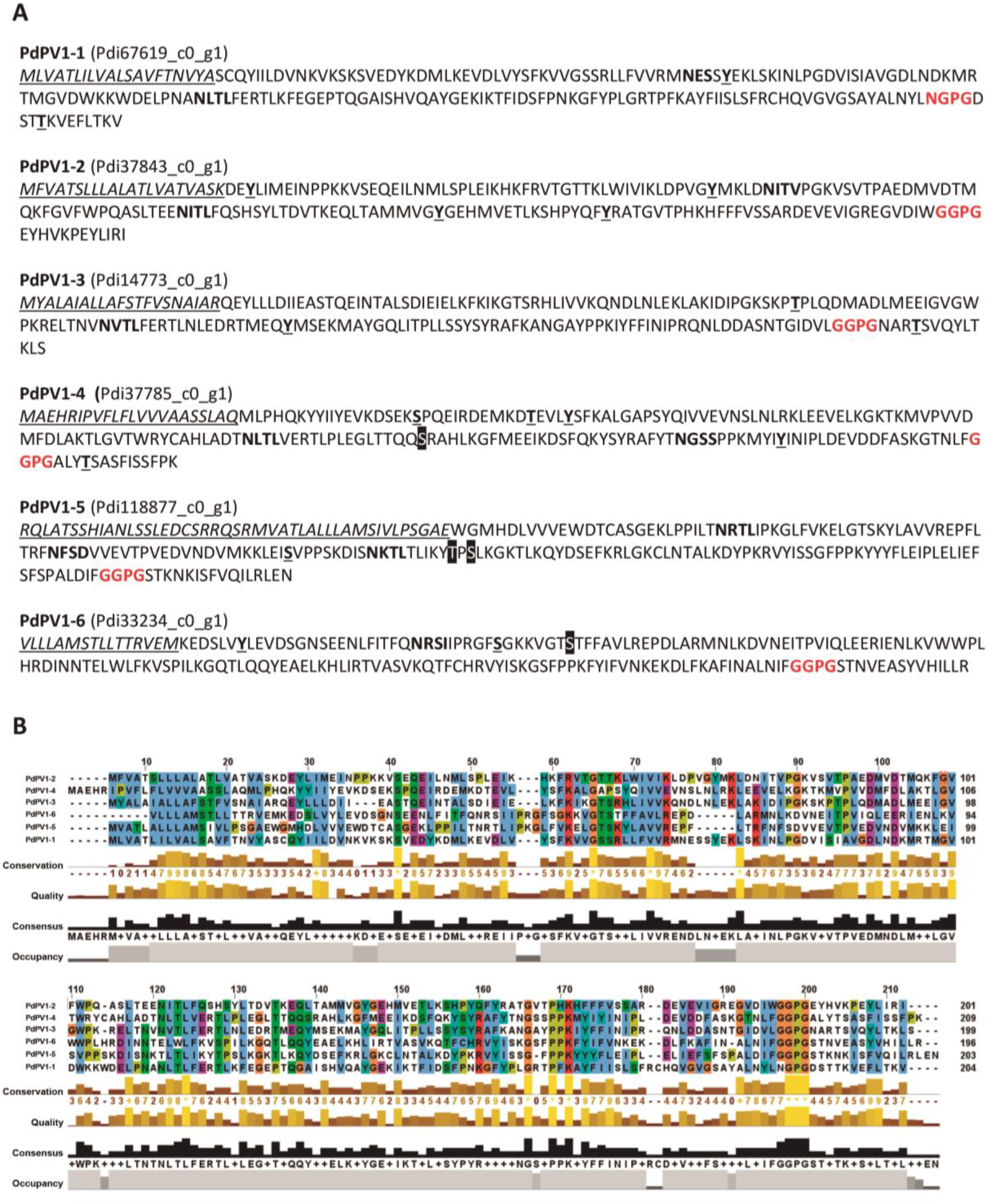
Subunit sequences of PdPV1. A) Deduced aminoacid sequences of the six PdPV1 subunits. Putative signal sequences are in italics and underlined. Potential phosphorilation sites are in bold underlined, potential N- and O-glycosylation sites are in bold and on black squares respectively. A conserved sequence is marked in bold red. B) Multiple sequence alignment of PdPV1 subunits.

